# Single-cell-resolution transcriptome map of human, chimpanzee, bonobo, and macaque brains

**DOI:** 10.1101/764936

**Authors:** Ekaterina Khrameeva, Ilia Kurochkin, Dingding Han, Patricia Guijarro, Sabina Kanton, Malgorzata Santel, Zhengzong Qian, Shen Rong, Pavel Mazin, Matvei Bulat, Olga Efimova, Anna Tkachev, Song Guo, Chet C. Sherwood, J. Gray Camp, Svante Paabo, Barbara Treutlein, Philipp Khaitovich

**Author notes:** Contributed equally (Ekaterina Khrameeva, Ilia Kurochkin, Dingding Han).

## Abstract

Identification of gene expression traits unique to the human brain sheds light on the mechanisms of human cognition. Here we searched for gene expression traits separating humans from other primates by analyzing 88,047 cell nuclei and 422 tissue samples representing 33 brain regions of humans, chimpanzees, bonobos, and macaques. We show that gene expression evolves rapidly within cell types, with more than two-thirds of cell type-specific differences not detected using conventional RNA sequencing of tissue samples. Neurons tend to evolve faster in all hominids, but non-neuronal cell types, such as astrocytes and oligodendrocyte progenitors, show more differences on the human lineage, including alterations of spatial distribution across neocortical layers.

## INTRODUCTION

The human brain is an enormously complex organ that has expanded greatly in comparison with the brains of our closest relatives, chimpanzees, bonobos, and other apes. Increased size alone, however, fails to explain cognitive abilities unique to humans [Semendeferi and Damasio, 2000; Elston et al., 2006; Teyssandier, 2008; Semendeferi et al., 2011; Barger et al., 2012]. Functional changes acquired on the human lineage are likely to be mediated by alterations in gene expression and cell type composition in particular brain structures [O’Bleness et al., 2012; Sousa et al., 2017; McKenzie et al., 2018]. Yet we currently lack a comprehensive understanding of these uniquely human evolutionary differences.

Gene expression within the human brain differs substantially among regions and anatomical structures, both within neocortex and among subcortical areas [Kang et al. 2011; Hawrylycz et al., 2012]. Initial studies comparing gene expression in humans to non-human primates (NHPs) examined one or several brain regions with the main focus on the prefrontal area of the neocortex [Enard et al., 2002; Caceres et al., 2003; Marvanová et al., 2003; Khaitovich et al., 2004b]. These studies identified multiple expression differences specific to humans and revealed an acceleration of expression evolution on the human lineage. While the expression differences shared among brain regions often represent molecular and functional changes not specific to the brain [Khaitovich et al., 2004a; Khaitovich et al., 2005], differences particular to individual brain regions tend to be associated with specific brain functions [Khaitovich et al., 2004a]. Recent studies examining eight and 16 brain regions in humans and closely related NHPs expanded these results further by revealing the rapid expression evolution of several subcortical regions in addition to the neocortical areas [Sousa et al., 2017; Xu et al., 2018].

While the brain is composed of functionally diverse anatomical regions, each brain region is composed of multiple cell types [Lein et al., 2007]. Single cell RNA sequencing provides an opportunity to decompose gene expression within brain regions and compare homologous cell types across species [La Manno et al., 2016; Saunders et al., 2018; Tosches et al., 2018; Zeisel et al., 2018]. A single cell level comparison between human and macaque transcriptomes in prenatal and adult dorsolateral prefrontal cortices indeed showed that all detected human cell types had a close homolog in macaques, and vice versa [Zhu et al., 2018]. Furthermore, this study identified genes differentially expressed between humans and macaques in individual cell types. Similar results were obtained in studies of single cell expression in human and chimpanzee cerebral organoids, indicating that cell type composition can be accurately matched between closely related primate species and expression differences within each type identified [Mora-Bermúdez et al., 2016; Pollen et al., 2019].

Here, we report transcriptome maps of the human, chimpanzee, bonobo, and macaque brain constructed using conventional RNA sequencing (bulk RNA-seq) and single nuclei sequencing (snRNA-seq).

## RESULTS

### Global gene expression variation analysis

We used bulk RNA-seq to examine RNA expression in 33 brain regions from four humans, three chimpanzees, three bonobos, and three rhesus monkeys (Fig. 1A,B; Table S1). The visualization of all expression differences revealed separation of species, consistent with their phylogenetic relationship (Fig. 1C,D; Fig. S1), as well as a common pattern of differences among the 33 brain regions within each brain (Fig. 1E,F; Fig. S1). Accordingly, the 33 brain regions further separated into seven clusters shared across individuals and species (Fig. 1G; Fig. S2). The clusters largely corresponded to anatomical areas, with three clusters representing cortical regions: primary and secondary cortices (cluster I), limbic and association cortices (cluster II), and archicortex (cluster III), while the remaining four clusters contained subcortical structures: thalamus and hypothalamus (cluster IV), white matter (cluster V), cerebellar grey matter (cluster VI), and striatum (cluster VII). This clustering was consistent with the one reported in humans based on gene expression analysis of 120 brain regions (Fig. S3) [Hawrylycz et al., 2012].

**Fig. 1.**
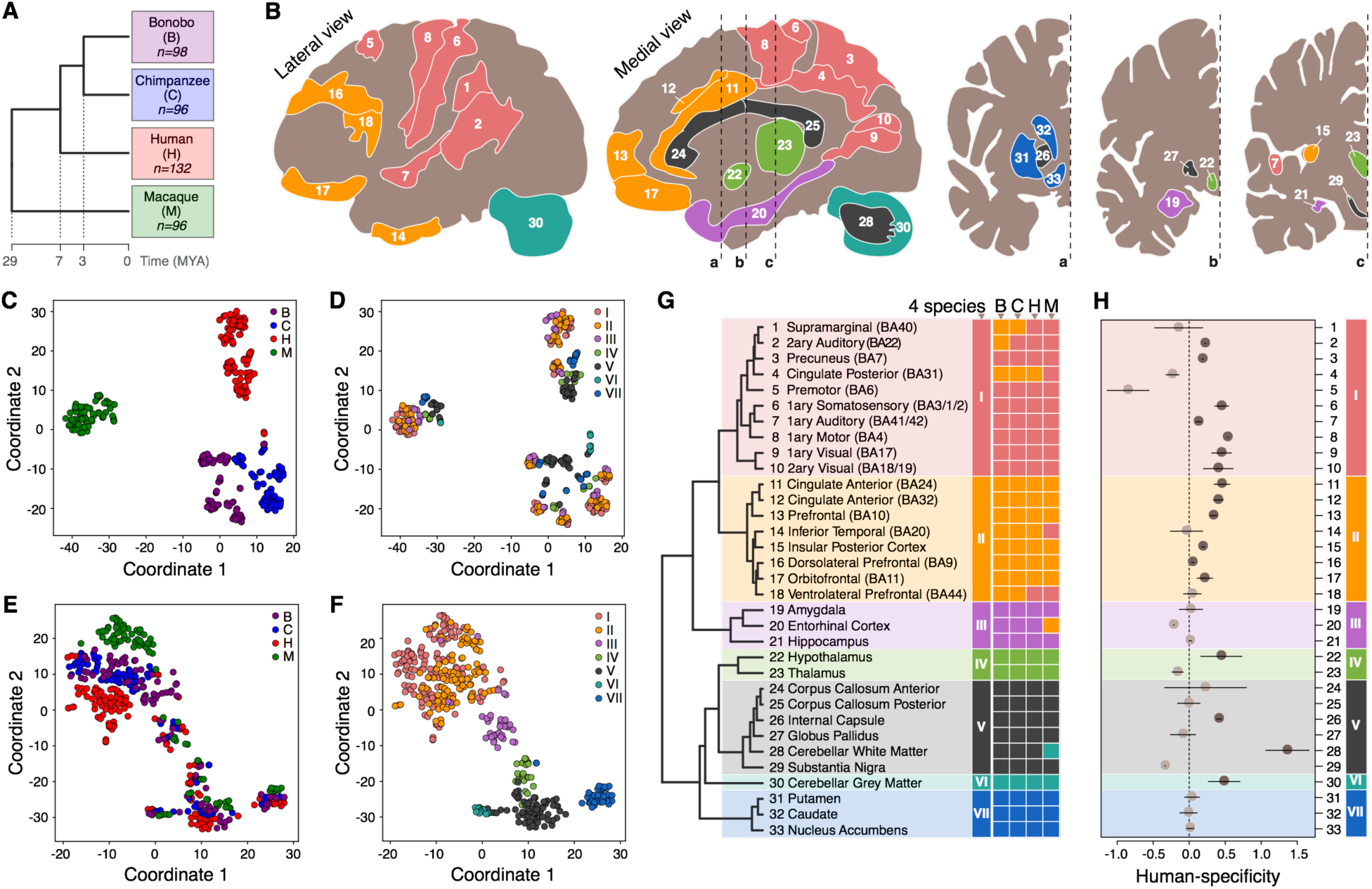
Gene expression variation analysis in 33 brain regions. **(A**) Phylogenetic relationship among analyzed species. Numbers indicate the number of analyzed brain samples. (**B**) Anatomical localization of 33 analyzed brain regions within the human brain. Colors represent expression-based regional clusters, defined in panel **G**. (**C**-**F**) tSNE plots based on expression variation among all 422 analyzed samples: the total variation (**C** and **D)**, and the residual variation after removal of the average species’ and individual differences (**E** and **F**). Each circle represents a sample. Circle colors represent species (**C** and **E**) or expression-based regional clusters (**D** and **F**). (**G**) Unsupervised hierarchical clustering of brain regions based on the average gene expression values of all 11,176 detected genes in four species. Regions within each species are assigned to the nearest cluster. The clustering based on each individual sample is shown in Fig. S2. (**H**) The human-specificity ratio of gene expression estimated in each brain region as the ratio of human-specific expression differences and chimpanzee-specific or bonobo-specific expression differences. Circles show the mean of chimpanzee-based and bonobo-based comparisons, and lines span the difference between the two estimates. Darker circles mark brain regions showing an excess of human-specific expression differences compared to both ape species.

Of the 11,176 detected orthologous protein-coding genes, 2,801 showed brain region-dependent species differences (ANOVA, FDR corrected p<0.00002; Fig. S4). Assigning differences among species to the evolutionary lineages recovered the phylogenetic relationship among the four species (Fig. S4,S5). Notably, region-dependent expression differences accumulated faster in the three cortical clusters compared to the subcortical regions across the phylogenetic tree (Fig. S4,S6).

### Regional analysis of the human-specific gene expression differences

Among 2,801 expression differences, we assigned the ones distinguishing humans from non-human primates (NHPs) to the human evolutionary lineage (Fig. S7, Table S2). The distribution of these differences was not uniform across brain regions. Neocortical regions represented in clusters I and II showed higher than the average proportion of human-specific expression differences, while the highest number of differences (1,079 genes) was found in cerebellar white matter (Fig. S8; Table S2). Normalizing the number of human-specific differences by the number of chimpanzee-specific or bonobo-specific ones (human-specificity ratio) revealed a similar pattern. The majority of neocortical regions in clusters I and II (72%), as well as the hypothalamus, internal capsule, and cerebellar white and gray matter showed an excess of the human-specific differences (Fig. 1H). Overall, in agreement with previous works [Sousa et al., 2017a; Xu et al., 2018], more differences mapped to the human lineage (the average n=712) than to the chimpanzee (n=641) or bonobo (n=640) lineages.

To test whether brain regions share human-specific expression differences, we first focused on the differences detected in more than 10 of the 33 brain regions (the average of human-chimpanzee and human-bonobo comparisons; Table S3). The 756 genes satisfying this criterion showed human-specific expression both in cortical and subcortical regions and were enriched in synaptic transmission, regulation of exocytosis, and neurotransmitter secretion terms (hypergeometric test, BH corrected p < 0.05, Figure S9). By contrast, genes showing human-specific expression in less than six brain regions did not display functional enrichment (hypergeometric test, BH corrected p > 0.05, n=655).

We further compared the relative expression human-specificity of brain regions determined in our study with previous reports. Despite the experimental and statistical differences of analyses, our results correlated positively and significantly with the published data [Sousa et al., 2017] (Spearman correlation, p < 0.01; Figure S10).

### Single nuclei transcriptome analysis

To investigate evolutionarily differences accumulating within cell types we sequenced RNA from individual nuclei (snRNA-seq) in three of the 33 brain regions of four species, anterior cingulate cortex (AC), caudate nucleus (CN), and cerebellar gray matter (CB), in three individuals per species (Table S1). To reduce experimental variation among species, tissue samples from one individual of each species were pooled and processed in parallel in each brain region (Fig. 2A). The nuclei species’ identity was then recovered computationally, based on the sequence differences between species. Each pool, except one (AC1), yielded two independent snRNA-seq libraries, resulting in a total of 88,047 nuclei with at least 500 unique detected molecules: 7,337±5,548 nuclei per species per region. Of them, 21% were derived from AC, 29% from CN and 50% from CB. Within each brain region, the humans were on average represented by 37% of the nuclei, chimpanzees by 10%, bonobos by 32%, and macaques by 21% (Table S4). Visualization of the total expression variation across these nuclei revealed a separation of the three brain regions (Fig. 2B; Fig. S11). The average expression levels of the nuclei within each tissue of each species correlated well with the corresponding bulk RNA-seq data and published single-cell RNA-seq data (Fig. 2C; Fig. S12,S13) [Pollen et al., 2019]. Similarly, the extent of human-specific expression divergence relative to the chimpanzee-specific or bonobo-specific divergence agreed well between the averaged snRNA-seq and the bulk RNA-seq data (Fig. 2D; Fig. S14). Within each region of each species, the nuclei formed six main clusters in AC and CN and four in CB, each enriched in known cell type markers (Fig. 2E-G; Fig. S15; Table S5). For purposes of our analysis, we focused on these broad cell classifications, but we are aware that subtypes could be more finely resolved and characterized (Fig. S16).

**Fig. 2.**
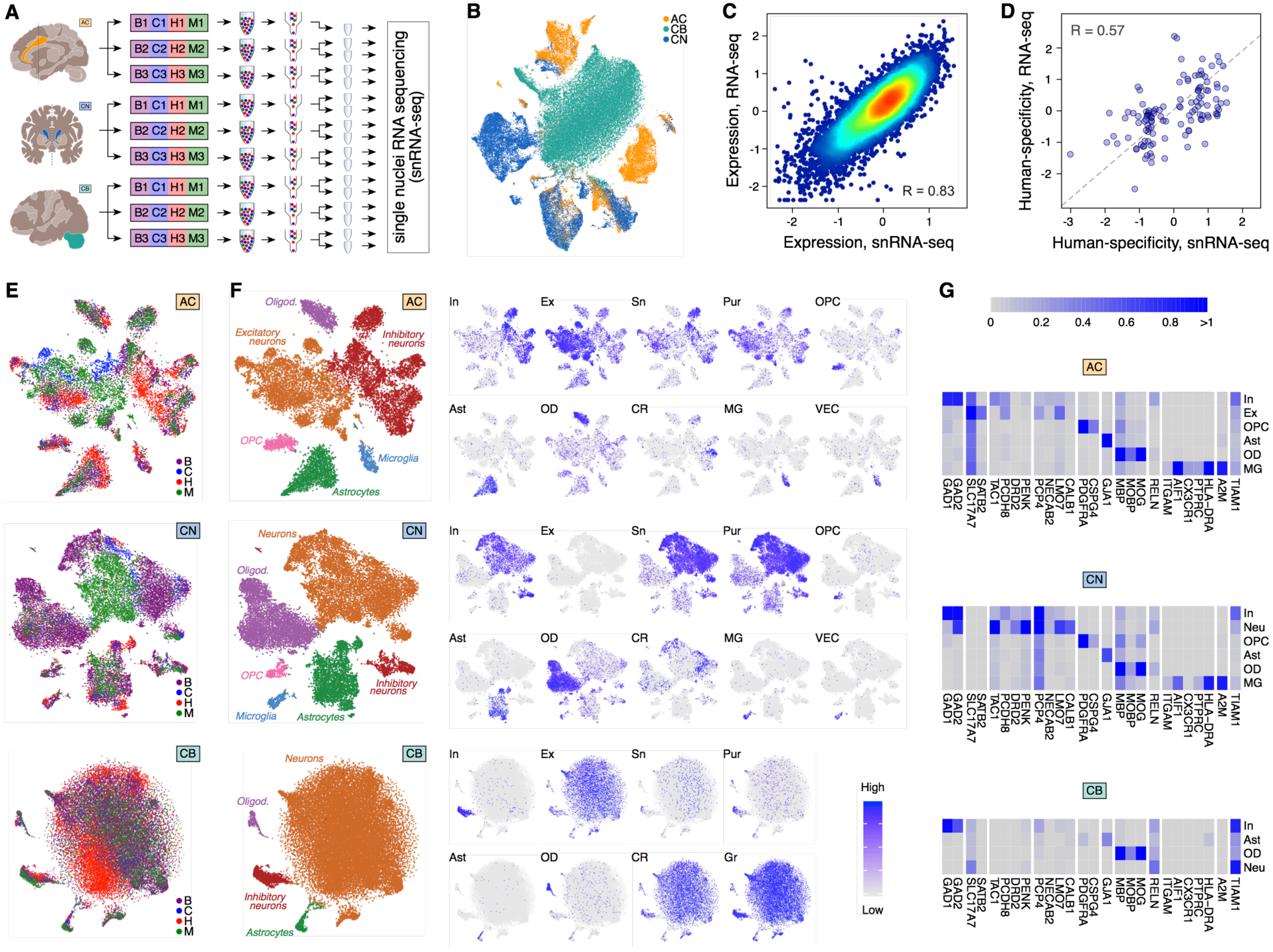
Single nuclei transcriptomics in three brain regions. **(A**) Design of the snRNA-seq experiment. (**B**) tSNE plot of 88,047 single nuclei colored by brain regions after integration with Seurat v3 [Stuart et al., 2019]. (**C**) Correlation of gene expression levels between bulk RNA-seq and averaged snRNA-seq datasets in human AC. Dots represent genes, and colors show the density of the dots. (**D**) Correlation of human-specificity ratios between bulk RNA-seq and averaged snRNA-seq datasets in AC for genes passing human-chimpanzee difference cutoff in either dataset. Each dot represents a gene. (**E**) tSNE plot of nuclei colored by species in each of the three brain regions after integration with Seurat v3 [Stuart et al., 2019]. (**F**) The cumulative cell type annotation of tSNE clusters (left) and projection of expression levels averaged across cell type marker genes onto the t-SNE plots. Abbreviations next to tSNE plots mark cell types: In – inhibitory neurons; Ex – excitatory neurons; Sn – spindle neurons; Pur – Purkinje cells; OPC – oligodendrocyte progenitor cells; Ast – astrocytes; OD – oligodendrocytes; CR – Cajal-Retzius cells; MG – microglia; VEC – vascular endothelial cells. (**G**) Average expression levels of cell type marker genes in tSNE clusters.

### Cell type-based analysis of human expression evolution in three brain regions

The calculation of the expression evolution velocity within each cell type across the human, chimpanzee, and bonobo lineages revealed that all neuronal subtypes, except cerebellar neurons, evolved at a faster rate than the rest of cell types (Fig. 3A,B). Of note, this result might be partially explained by the larger cell type heterogeneity of neuronal clusters (Fig. S17).

**Fig. 3.**
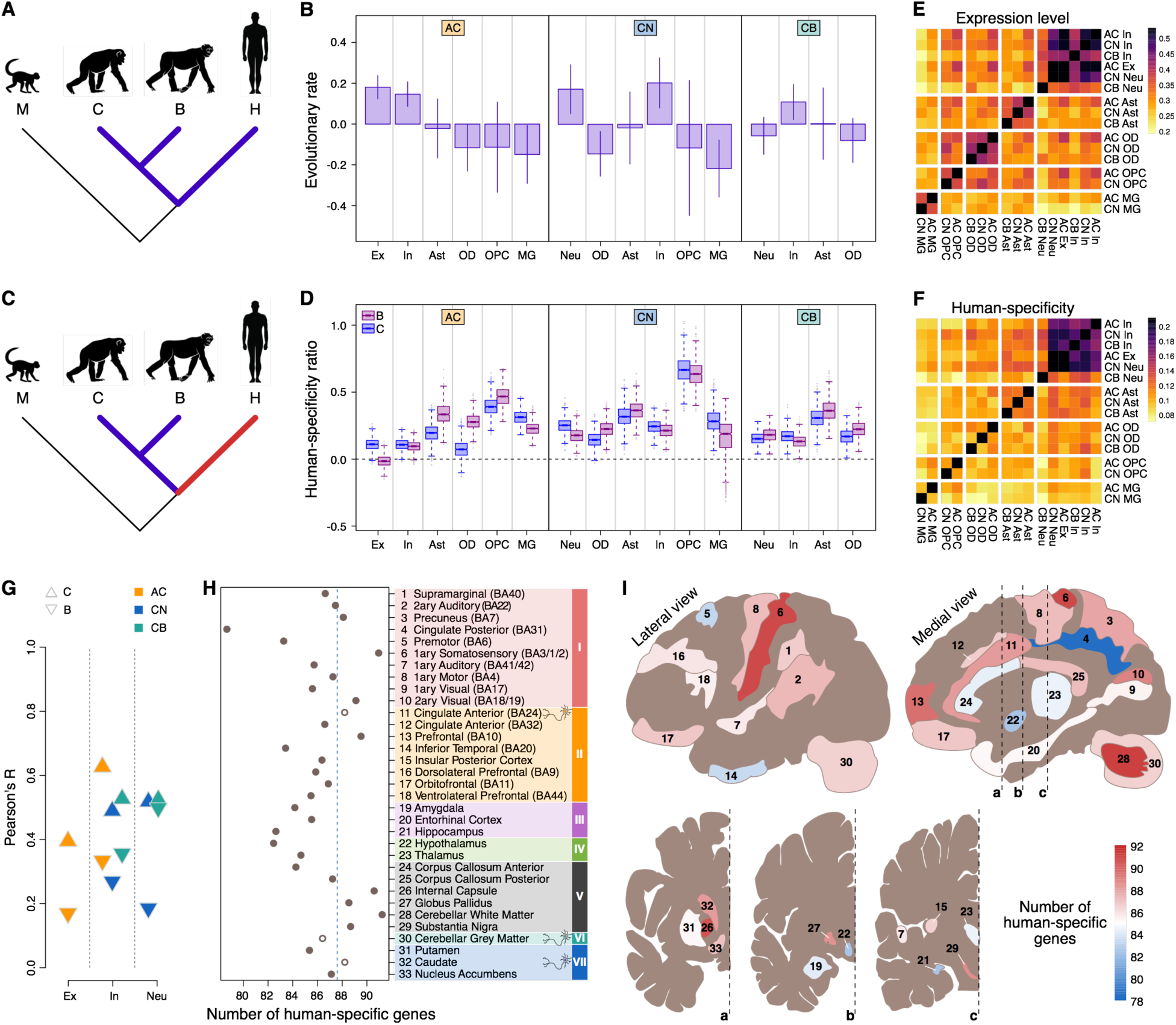
Cell-type-based analysis of the expression evolution in three brain regions. **(A**) Phylogenetic tree highlighting the branches used in the evolutionary velocity analysis. (**B**) The evolutionary velocity of cell types within each brain region normalized by the mean of the region. Error bars mark the standard deviation of the average estimates. (**C**) Phylogenetic tree highlighting the branches used in the human-specificity ratio analysis. (**D**) Human-specificity ratio calculated within each tSNE cluster in each of the three brain regions. The ratio represents the number of genes with human-specific expression divided by the number of genes with chimpanzee-specific (blue boxplots) or bonobo-specific (purple boxplots) expression. Boxes mark the median and the first and the third quartiles of the distribution, and whiskers extend to the 1.5 interquartile ranges. The cell types are abbreviated as in Fig. 2F. (**E**) The expression level similarity among tSNE clusters based on the average gene expression levels within clusters in humans. The colors here and in panel F indicate Pearson correlation coefficients. (**F**) The similarity of human-specific ratio estimates among tSNE clusters calculated based on the comparison to chimpanzee and bonobo combined. (**G**) Correlation of human-specificity ratios defined as human-macaque difference relative to chimpanzee-macaque (triangles) or bonobo-macaque difference (inverted triangles) between bulk RNA-seq and snRNA-seq datasets for genes preferentially expressed in a specific cell type (Table S5). Colors indicate brain regions. X-axis labels indicate neuronal subtypes. (**H**) Numbers of genes showing human-specific expression in each brain region in bulk RNA-seq dataset overlapping with genes showing human-specific expression in at least one of the neuronal subtypes in snRNA-seq data. Empty circles indicate the three brain regions used in snRNA-seq experiment. (**I**) Anatomical localization of 33 regions in the human brain colored according to the overlap between human-specific expression differences in bulk RNA-seq and in neuronal subtypes.

Remarkably, the comparison of the expression evolution velocities between the human lineage and the chimpanzee or bonobo lineages revealed an opposite picture (Fig. 3C,D). All neuronal clusters from all brain regions, including the cluster of cerebellar granular cells, showed smaller excess of human-specific expression differences over the chimpanzee- or bonobo-specific ones, than the other cell types (Fig. 3D). Specifically, astrocytes and oligodendrocyte progenitor cells consistently showed the largest excess of human-specific expression differences among examined cell types (Fig. 3D). Notably, the human-to-NHPs evolutionary ratio estimates calculated independently using chimpanzee or bonobo data were highly consistent (Fig. 3D).

Gene expression within each cell type correlated well across brain regions (Fig. 3E). By contrast, human-specific expression differences correlated well among neuronal subtypes excluding cerebellar granule cells, but not among astrocytes, oligodendrocytes, oligodendrocyte precursors, or microglia originating from different brain regions (Fig. 3F). This result indicates that most of neurons, but not the other analyzed cell types, share a characteristic signature of human-specific expression differences particular to neurons.

### Deconvolution of bulk human-specific expression differences using neuronal evolutionary signature

The existence of the neuronal evolutionary signature shared among brain regions could be used to deconvolute bulk RNA-seq data, analogous to the cell type marker-based deconvolution procedure [Wang et al., 2019]. Supporting this notion, human-specificity ratios of genes preferentially expressed in neuronal subtypes correlated positively and significantly between single nuclei and bulk RNA-seq datasets (Fig. 3G).

To estimate the extent of human-specific neuronal differences in each of the 33 brain regions using bulk RNA-seq data, we calculated the overlap between genes showing human-specific expression differences in bulk RNA-seq data and in neuronal subtypes. Because this overlap should be positively biased towards the three brain regions used in both bulk and single nuclei analysis, most of the brain regions contained fewer neuronal human-specific differences compared to them (Fig. 3H). Yet, seven brain regions, including primary somatosensory cortex, internal capsule, and cerebellar white matter containing some of the deep nuclei of the cerebellum, displayed more differences, suggesting extensive human-specific alterations of neuronal expression in these regions (Fig. 3H,I).

### Gene expression differences detected by snRNA-seq and bulk RNA-seq

Notably, up to 12% of the differences separating humans from apes detected using bulk RNA-seq were also present in snRNA data (> 2-fold difference in Homo/Pan comparison, BH-adjusted p < 0.05, two-sided t-test for RNA-seq, Wilcoxon test implemented in *seurat* for snRNA-seq; Fig. 4A,B). Furthermore, the expression differences detected in multiple cell types showed a higher difference amplitude in bulk RNA-seq compared to the differences detected in particular cell types (Fig. 4C). Since 93-94% of genes expressed in the brain are detected in multiple cell types, the differences present in just one cell type might be attenuated or lost in the bulk tissue samples.

**Fig. 4.**
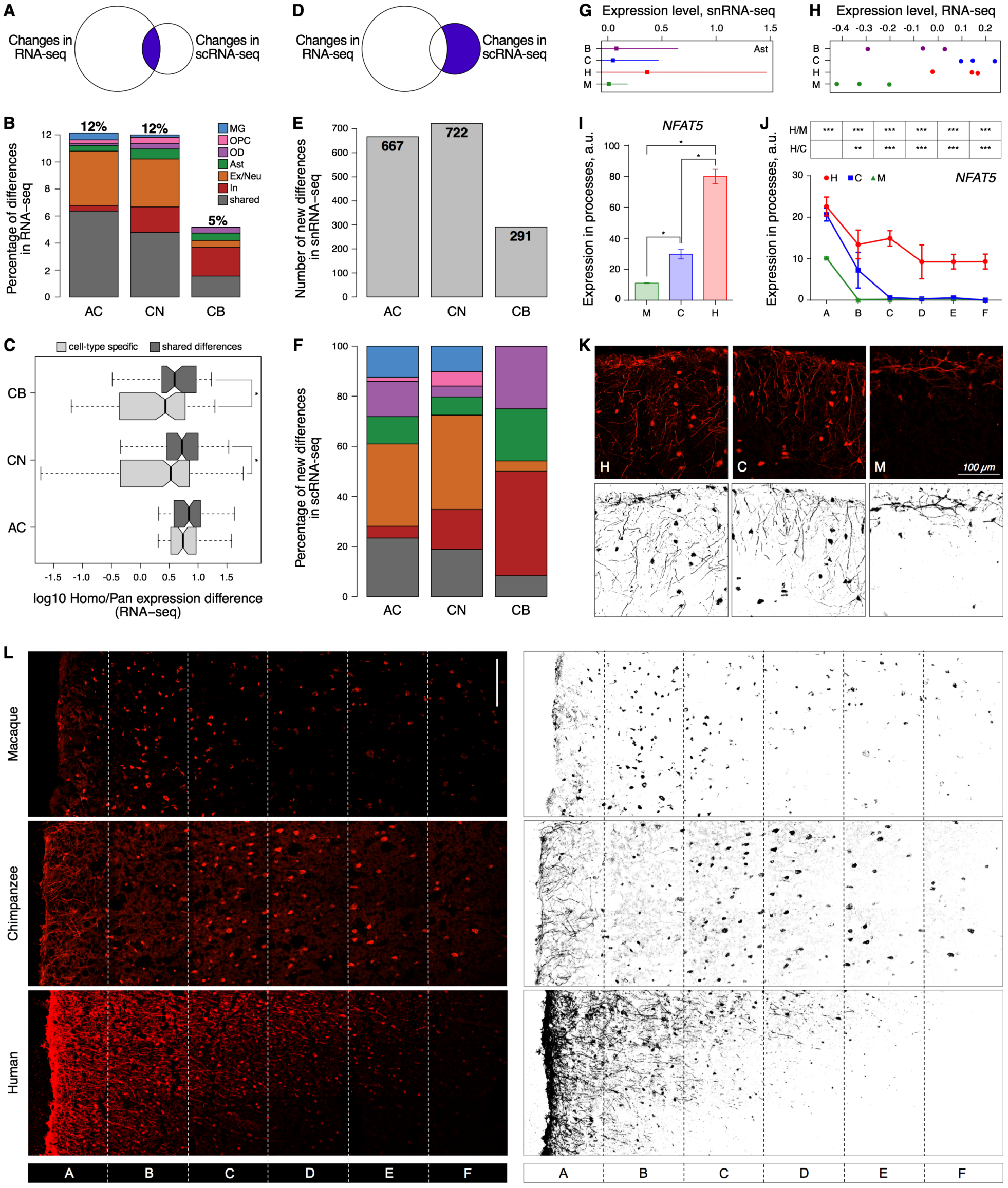
Gene expression differences detected by snRNA-seq, bulk RNA-seq, and IHC. **(A)** Schematic representation and **(B)** Percentage and the cell type specificity of the expression differences present in bulk RNA-seq and snRNA-seq. **(C)** The amplitude of expression differences detected in one (specific) or multiple (shared) cell types in bulk RNA-seq. * – p<0.05 (one-sided t-test). **(D)** Schematic representation, (**E**) Numbers, and **(F)** Cell type specificity of the expression differences solely detected by snRNA-seq. Color as in panel A. **(G)** The mean log10-transformed expression level of *NFAT5* mRNA in AC astrocytes (squares), and the standard deviation of the mean (vertical lines). **(H)** The log10-transformed read counts normalized for the median of *NFAT5* mRNA in bulk AC data. Circles indicate individual samples. **(I)** Average fluorescent intensities of NFAT5 IHC signal in the astrocytic processes of macaques, chimpanzees, and humans across cortical layers and (**J**) at different cortical depth. Error bars show the standard deviation of the mean. *** – p<0.0005, ** – p<0.0005, * – p<0.05 (two-sided t-test, Holm-Sidak correction). H/C – human-chimpanzee comparison. H/M – human-macaque comparison. Symbols indicate cortical sections located at increasing depth, depicted in panel L. **(K)** IHC (upper panels) and its binarized representation (lower panels) of NFAT5 protein in the uppermost layer of AC sections. (**L**) Immunostaining (left panels) and its binarized representation (right panels) of NFAT5 protein in the three upper layers of AC sections in macaques, chimpanzees, and humans (see also Figures S20-S23 for GFAP and DAPI staining of these sections). Sections A-F indicate segmentation areas used in the analysis presented in panel J. Scale bar 100 µm.

Indeed, in AC we identified 667 genes showing the expression differences separating humans from apes using snRNA-seq, not detectable in bulk RNA-seq data (Fig. 4D-F). Similarly, there were 722 such differences in CN, and 291 in CB. On average, these differences constitute more than two-thirds of the differences found using snRNA-seq.

Among the 667 expression differences found in AC, 32.8% localized in excitatory neurons, and only 4.7% – in inhibitory neurons. Notably, the differences particular to excitatory neurons were enriched in synaptic transmission terms, while the differences detected in microglia (12.5%) were enriched in immune response and lipid localization functions (hypergeometric test, BH corrected p < 0.01; Fig. S18).

### Gene expression differences detected by immunohistochemistry

We visualized the spatial distribution of expression differences revealed by snRNA-seq in AC using immunohistochemistry (IHC). As neurons consistently showed the smallest excess of human-specific expression differences in snRNA-seq data, we focused on the other cell types for IGH analysis. Specifically, we selected two genes, *MSI2* and *NFAT5*, which displayed higher expression in human astrocytes according to snRNA-seq data (two-sided t-test, p-value<10^−10^; Fig. 4G; Fig. S19). Both genes generated clear and specific staining in the frozen AC tissue sections. Of note, while *MSI2* similarly showed an increased expression in human AC in conventional RNA-seq data (two-sided t-test, p=0.0001; Fig. S19), there was no such expression increase for *NFAT5* (two-sided t-test, p=0.5; Fig. 4H). Immunohistochemical staining assisted by the common astrocytic and neuronal marker proteins, GFAP and NeuN, placed MSI2 and NFAT5 proteins within astrocytic processes and neuronal cell bodies in human, chimpanzee, and macaque tissue sections (Fig. S20-S23). Remarkably, both proteins demonstrated significantly higher fluorescent intensity in human astrocytic processes compared to those in NHPs (two-sided t-test, p < 0.05; Fig. 4I). Furthermore, while the processes localized in the uppermost cortical layer in chimpanzees and macaques, they spread to deeper laminar structures, including layers two and three, in humans (two-sided t-test, p < 0.05; Fig. 4J,K; Fig. S24). This observation adds to reports of functional heterogeneity and rapid evolution of astrocytic cell types in primates [Oberheim et al., 2009; Haim and Rowitch, 2017].

## DISCUSSION

We show that the distribution of human-specific expression differences separating us from our closest living relatives, chimpanzees and bonobos, was not uniform across 33 examined brain regions resembling functional networks defined by magnetic resonance imaging studies (Fig. S25). Instead, our results suggest an intricate pattern of the expression evolution of human brain involving both neocortical and subcortical regions.

Our analysis of human-specific expression features conducted at the single nuclei level further provided the following insights:

First, we detected multiple expression differences between species within each cell type. While approximately one third of these differences, especially the ones present in multiple cell types, could be detected using conventional bulk RNA-seq data, the remaining differences could only be revealed using cell-type-specific methodologies.

Second, the majority of neuronal subtypes evolved faster on the human and ape lineages, then the other cell types. This effect might be partially explained by greater heterogeneity of broadly defined neuronal cell type clusters [Zeisel et al., 2018].

Third, non-neuronal cell types showed substantially greater excess of human-specific expression differences in all three examined brain regions compared to neurons. Among them, astrocytes and oligodendrocyte progenitors displayed the largest excess of human-specific expression differences. These human-specific differences were particular to each brain region, precluding deconvolution of bulk RNA-seq data for non-neuronal cell types.

It has to be noted that the depth of our study was not sufficient to accurately examine expression evolution in specific neuronal subtypes, such as von Economo neurons, known to be overrepresented in human AC and the frontoinsular cortex [Yang et al., 2019], or the rosehip neurons [Boldog et al., 2018]. Thus, our results do not exclude the existence of specialized neuronal populations showing rapid expression evolution in humans. Furthermore, application of neuronal human-specific expression signature detected in snRNA-seq data to 33 brain regions revealed seven, including primary somatosensory cortex, internal capsule, and cerebellar white matter, showing the greater extent of human-specific expression differences characteristic of neuronal cells.

While all reported cell-type-specific evolutionally differences are novel, they concur with previous observations. The regional specificity of the astrocytic human-specific expression differences matches reports of molecular and functional heterogeneity of astrocytes in adult brain regions [Haim and Rowitch, 2017]. In turn, excess of astrocytic human-specific expression differences matches histological differences reported between humans and the other primate species for interlaminar astrocytes, polarized astrocytes, and varicose projection astrocytes [Oberheim et al., 2009; Falcone et al., 2018]. Similarly, oligodendrocyte progenitor cells showing rapid expression evolution in human caudate nucleus were reported to dysfunction in the caudate nucleus of schizophrenia patients [Georgieva et al., 2006; Uranova et al., 2007; Cassoli et al., 2015; Mauney et al., 2015], a disorder suggested to affect aspects of cognition particular to humans [Konopka and Geschwind, 2010; Dean, 2009].

Taken together, our results show that systematic investigation of gene expression evolution across a large number of brain regions and cell types has the potential to reveal evolutionary patterns reflecting the emergence of the human brain functionality. An important component that was missing from our study, an analysis of temporal patterns of expression evolution in the developing brain, analogous to the one presented in [Zhu et al., 2018], would further increase the power to associate expression differences with cognitive functions.

## METHODS

### Samples

Human samples were obtained from the Chinese Brain Bank Center. Chimpanzee samples were obtained from the Southwest National Primate Research Center in San Antonio, Texas. Bonobo samples were obtained from the Lola Ya Bonobo Sanctuary, Democratic Republic Congo. Rhesus monkey samples were obtained from the Suzhou Experimental Animal Center, China.

This study was reviewed and approved by the Institutional Animal Care and Use Ethics Committee at the Shanghai Institute for Biological Sciences, CAS. Informed consent for the use of human tissues for research was obtained in writing from all donors or their next of kin. All non-human primates used in this study suffered sudden deaths for reasons other than their participation in this study and without any relation to the tissue used.

### Sample dissection

A total of 422 brain samples were dissected from the brains of 14 individuals with at least 3 individuals per species (Table S1). For each individual, samples were dissected from 33 brain regions covering all major anatomical and functional brain structures (Table S1). All brains were previously frozen in liquid nitrogen (humans, chimpanzees, macaques) or in isopentane/dry ice (bonobos) and stored at −80°C until used. All human brains, one chimpanzee brain and one macaque brain were sliced as separate hemispheres in coronal orientation before freezing and storage; the remaining brains were frozen and stored as entire hemispheres and, before sample dissection, hemispheres were atemperated to −15°C and sliced in coronal orientation. All brain slices were stored at −80°C. For sample dissection, *The Atlas of the Human Brain* [Mai et al., 2016] and *The Rhesus Monkey Brain* [Paxinos et al., 2009] were used to locate the areas of interest in human and macaque brains respectively. As there is no equivalent published resource for chimpanzee and bonobo brains, chimpanzee and bonobo areas were located using *The Atlas of the Human Brain*. Hemispheres/slices were atemperated to −20°C prior to dissection, placed on a metal board previously cleaned with 75% ethanol and chilled at −80 C, and pieces of 10-60 mg were cut out of selected areas using a metal scalpel. Dissected samples were then collected with tweezers, put into tubes and immediately stored at −80°C. Dissection was performed on dry ice. All materials used during dissection (scalpels, tweezers, tubes) were sterile and chilled in dry ice at −80°C before use.

### RNA Sequencing (RNA-seq)

Total RNA was isolated using Direct-zol™96 RNA (ZYMO RESEARCH,) from 10 mg of the frozen tissue per sample. Sequencing libraries were prepared with NEBNext® Ultra™ II RNA Library Prep Kit (New England Biolabs) according to manufacturer’s instructions. Briefly, poly-T oligo-attached magnetic beads were used to isolate long polyadenylated RNA from 100 ng of total RNA. After fragmentation, first-strand cDNA was reverse transcribed with random hexamer-primers, followed by second-strand cDNA synthesis, end repair, adenylation of 3’ ends, and ligation of the adapters. Fragments were then enriched by PCR and sequenced on the Illumina HiSeq 4000 system using the 150-bp paired-end sequencing protocol. All samples were randomized with respect to species prior to library preparation and RNA sequencing.

### Single-nuclei sequencing (snRNA-seq)

Frozen tissue samples of cingulate anterior cortex (BA24), cerebellar gray matter, and caudate nucleus from brains of three individuals per species were used for the intact nuclei isolation. For each brain region, three pooled sample sets were prepared. Each sample set contained pooled equal tissue samples of 5 mg for one human, one chimpanzee, one bonobo, and one macaque individual (Fig. 2A).

All steps were performed on ice, and spinning of the samples was performed at 4°C. The pooled tissue pieces for each set were minced on ice using a scalpel and then washed into a Dounce homogenizer using PBSE (PBS + 2 mM EDTA) + 1% BSA + 0.3 M Sucrose and dounced with 10 strokes using pestle A followed by 10 strokes using pestle B. The homogenate was transferred to a 15 ml tube and spun down 5 min at 900xg. The supernatant was aspirated, the pellet resuspended 20 times in PBSE + 1% NP-40 and incubated for 7 min on ice to deliberate nuclei. The homogenate was spun down 5 min at 900xg, and the supernatant was aspirated. The pellet was washed 2 times using PBSE + 1% BSA and once using PBS + 1% BSA. Nuclei were stained using DAPI (BD Pharmingen, 1:1000 dilution) and filtered through a 30 um strainer (Miltenyi Biotec) before sorting. Sorting was performed using a FACS Fusion (BD) to sort for DAPI+ positive events and to remove debris and doublets. After sorting, nuclei were spun for 5 min at 900xg and resuspended in PBS + 1% BSA to be loaded on the 10x microfluidic chip device. All, except one (ACC1), of the obtained single nuclei suspensions were loaded on two lanes of a 10x microfluidic chip device.

Single nuclei experiments were performed using a 10x Chromium single cell 3’ v2 reagent kit by precisely following the detailed protocol of the manufacturer to construct 10x Genomics single-cell 3’ libraries. Each library was barcoded using the i7 barcodes provided by 10x. Single nucleus libraries were pooled at equal ratios and run using paired-end sequencing on the NovaSeq 6000 platform (Illumina) by following manufacturer’s instructions.

### Immunohistochemistry

For multiple immunofluorescent histochemistry, 20 µm thick cryosections were prepared from samples of the anterior cingulate cortex (BA24) from three humans, three chimpanzees, and three rhesus monkeys (Table S1). All samples were previously frozen in liquid nitrogen and stored at −80°C until used.

Sections were thaw-mounted onto microscope slides and fixed with 4% paraformaldehyde solution for 7 min followed by washing in phosphate-buffered saline – 0.1M PBS (pH 7.4) three times for 5 min. Then sections were preincubated in PBST (0.5% Triton X-100 in PBS) with 5% normal donkey serum (NDS) and 5% normal horse serum (NHS) for 1 h. Reaction with a mixture of primary antibodies (Table S6) consisting of one antibody against the human-specific antigen and one cell-type marker was performed in the blocking buffer (2.5% NDS and 2.5% NHS in PBST) for 24 h at 4°C in Shandon coverplates. We used rabbit anti-NFAT5 (NB120-3446) and mouse anti-MSI2 (NBP2-45837) antibodies against human-specific antigens, as well as glial goat anti-GFAP (PA5-18598), neuronal mouse anti-NeuN (MAB-377), and rabbit anti-NeuN (24307) antibodies against cell-type markers.

Following washing and incubation with biotinylated horse anti-rabbit (BA-1100) or mouse (BA-2000) IgG corresponding to primary human-specific antigen antibody, in the blocking buffer for 2 h at room temperature, sections were rinsed in PBST. Sections were processed with a mixture of donkey anti-goat Alexa Fluor 488 (A-11055) or anti-mouse Alexa Fluor 488 (A-21202) / anti-rabbit Alexa Fluor 488 (A-21206) and streptavidin-Alexa Fluor 568 conjugate (S-11226) for 2 h (Table S6). After a wash in PBST, sections were incubated in 1% Sudan black B solution in 70% ethanol for 10 min to block lipofuscin autofluorescence. Then washed in PBS and mounted with Fluoromount aqueous mounting medium (Sigma) with blue fluorescent nuclear counterstain DAPI, coverslipped and sealed with nail polish. No staining was seen in control sections processed without the primary antibody staining.

Images were obtained by Zeiss LSM 800 AiryScan system with C Plan-Apochromat 40x/1,3 Oil DIC UV-VIS-IR objective.

### RNA-seq data processing

In total, we obtained 7,483,498,084 RNA-seq reads, with an average sample coverage of approximately 17.7 million reads (Table S7). To remove Illumina universal adapters and low-quality bases at the ends of reads, we used trimmomatic [Bolger et al., 2014] with the following parameters: “PE -phred33 ILLUMINACLIP:all.fa:2:30:10:2:true SLIDINGWINDOW:4:15 LEADING:3 TRAILING:3 MINLEN:20”. The union of all adapter sequences provided by trimmomatic was used, as well as an additional sequence of Illumina universal adapter found by fastQC (AGATCGGAAGAG), for palindrome clipping. Reads were further mapped to the corresponding reference genomes (GRCh38, Mmul_8.0.1, panpan1.1 and Pan_tro_3.0) using hisat2 [Kim et al., 2015] with the following parameters: “--no-softclip --max-intronlen 1000000 -k 20”. Gene expression levels were estimated as Transcripts Per Million (TPM) using stringtie [Pertea et al., 2015] with the following parameters: “-e -G reference.gtf -B out -A out.tab”. Reference genome sequences, gene annotations and orthologous gene tables for all species were obtained from Ensembl v91 [Zerbino et al., 2018]. One-to-one orthologous protein-coding genes with TPM>1 were used in further analysis. TPMs were further normalized by the sample median, and log-transformed.

tSNE was applied to visualize the samples (Fig. 1B,E). To reduce individual-to-individual variability, gene expression values were additionally normalized by the median expression level among regions in each individual brain (Fig. 1C,F; Fig. S1).

Complete linkage method of unsupervised hierarchical clustering with one minus Pearson correlation coefficient as a distance metric was used to cluster brain regions based on average gene expression values among four species (Fig. 1G). Further, we calculated the average gene expression values within each cluster. Based on them, we assigned each brain region of each species to the nearest cluster using one minus Pearson correlation coefficient as a distance metric, to assess the conservation of clustering among four species (Fig. 1G; Fig. S2).

Additionally, to test the robustness of this clustering procedure, we compared our clusters with previously published data from Allen Human Brain Atlas (AHBA) [Hawrylycz et al., 2012]. First, we selected regions from AHBA that correspond to regions from our dataset. Next, we assigned a cluster label to each region from AHBA based on the association between the regions from AHBA and our dataset. Further, we calculated average gene expression values within each cluster of AHBA and our dataset. Finally, we calculated the pairwise Pearson correlation coefficient for each corresponding cluster between AHBA and our dataset (Fig. S3; Table S8).

### Expression differences between species

First, we compared gene expression levels that were obtained in our study with a previously published dataset containing 16 brain regions in human and non-human primates [Sousa et al., 2017a]. We employed the same read mapping and counting procedures as described above for RNA-seq reads from the National Center for Biotechnology Information BioProjects database, accession number PRJNA236446 [Sousa et al., 2017a]. The resulting TPM values were log2 transformed, and then quantile normalization was applied. The correspondence between the brain regions in [Sousa et al., 2017a] and in our dataset was based on anatomical localization of regions in the human brain. Separately for each dataset and region, we classified genes that demonstrated expression differences between human and chimpanzee using t-test (p-value < 0.05). For genes that passed t-test threshold in both datasets, we calculated log2-fold changes between gene expression levels in human and chimpanzee (Fig. S10A). To check if log2-fold change values were in agreement between datasets, we performed Fisher’s test (Fig. S10B). The same analysis was done for comparison of gene expression levels between human and macaque (Fig. S10C,D), and between human and average gene expression in chimpanzee and macaque (Fig. S10E,F).

To identify expression differences among species that varied significantly depending on the brain region, ANOVA was applied with both species and regions variables as factors (Fig. S4). To reconstruct the phylogenetic tree based on the identified expression differences, UPGMA method was used with one minus Pearson correlation coefficient as a distance metric (Fig. S4). Similarly, phylogenetic trees were reconstructed for each brain region separately (Fig. S5), and, for each tree, the total branch length was calculated (Fig. S6).

To assign the expression differences to the evolutionary lineages, we classified human-specific expression differences as the ones showing 2-fold greater human/macaque difference relative to chimpanzee/macaque or bonobo/macaque differences (Fig. S7,S8). Chimpanzee-specific expression differences were defined as the ones showing 2-fold greater chimpanzee/macaque difference relative to human/macaque difference. Bonobo-specific expression differences were defined similarly. The human-specificity ratio of each brain region was estimated as the number of human-specific genes divided by the number of chimpanzee-specific or bonobo-specific genes (Fig. 1H).

To cluster genes with expression differences among species that varied significantly depending on the brain region, UPGMA method was used with one minus Spearman correlation coefficient as a distance metric (Fig. S26). A minimal number of modules (n=3) were selected for further functional network analysis (Fig. S25): M1 (1,389 genes), M2 (1,267 genes), and M3 (145 genes).

### Single-nuclei data processing

A total of 3,081,653,593 paired-end sequencing reads of snRNA-seq were processed using publicly available 10x Genomics software – Cell Ranger v2.2.0 [Zheng et al., 2017]. At the first step, *cellranger mkfastq* was used to convert binary base call (BCL) files to FASTQ files and to decode the multiplexed samples simultaneously. Next, *cellranger count* was applied to the obtained FASTQ files. It performed sequencing alignment using STAR v2.5.3a to a concatenation of human (hg38), chimpanzee (panTro5), bonobo (panPan2) and macaque (rheMac8) reference genome assemblies.

To assign each nucleus to a species, we first used a custom Perl script to calculate the number of UMIs mapped to each species reference genome per nucleus, based on BAM files generated by cellranger. Another custom R script was used to assign a nucleus to species. First, the table was normalized for the total number of UMIs per species, to balance the mappability differences arising due to the evolutionary differences between the primate species. Then, a nucleus was assigned to a particular species if >50% of its UMIs were mapped to this species. The threshold of 50% was chosen based on the distribution of maximal proportions of UMI mapped to one species per nucleus (Fig. S27). A total of 107,019 nuclei assigned to species with at least 500 unique detected molecules were used in further analysis (Table S9).

To calculate gene expression values, we remapped each nucleus to the reference genome assembly of an assigned species using *cellranger count*. It performed sequencing alignment using STAR v2.5.3a to human (hg38), chimpanzee (panTro5), bonobo (panPan2) and macaque (rheMac8) reference genome assemblies separately. To generate the gene expression matrix, a list of UMIs in each gene and within each nucleus was assembled, then UMIs within ED = 1 were merged together. The total number of unique UMI sequences was counted, and this number was reported as the number of transcripts of that gene for a given nucleus. A total of 88,047 nuclei were reported by *cellranger count* at this step.

Additionally, to confirm that gene expression values were calculated correctly, we applied an alternative procedure of gene expression calculation and nucleus-to-species assignment based on the human-chimpanzee-bonobo-macaque consensus genome [Kanton et al., 2019], and obtained highly similar results (Fig. S28).

The sparse expression matrix generated by *cellranger* analysis pipeline with the list of 88,047 nuclei assigned to species was used as input to the Seurat software v3.0 [Stuart et al., 2019]. To account for technical variation, we performed cross-species integration. At the first step, for each species separately, we performed normalization using “LogNormalize” with the scale factor of 10,000 and identified 2,000 variable features. Next, we performed cross-species integration by finding corresponding anchors between the species using 30 dimensions. We then computed 50 principal components and tested their significances by JackStraw. We selected the first 30 principal components for subsequent tSNE and clustering analyses.

To compare snRNA-seq gene expression levels with RNA-seq dataset, we first calculated average gene expression values across all nuclei per region per species, then log-transformed, normalized by the median, and divided by the gene length to obtain RPKMs. In each species and region, Pearson correlation coefficients were calculated between snRNA-seq RPKMs and bulk RNA-seq RPKMs for all genes expressed in both datasets (Fig. 2C, Fig. S12). To calculate bulk RNA-seq RPKMs for this analysis, we normalized log-transformed read counts per gene by the sample median but did not normalize by the median expression level among regions in each individual brain, for consistency with snRNA-seq dataset. Additionally, we calculated Pearson correlation coefficients between snRNA-seq and RNA-seq human-specificity ratios for genes with dramatic (> 100 times) human/chimpanzee gene expression differences in either snRNA-seq or RNA-seq (Fig. 2D, Fig. S14).

We further compared the average snRNA-seq RPKMs with published single-cell RNA-seq dataset (Fig. S13) [Pollen et al., 2019]. For this analysis, we obtained average gene expression values for the primary brain samples and the organoid models from GEO accession GSE124299, log-transformed them, normalized for the sample median and divided by the gene length to obtain RPKMs.

*Seurat 3*.*0* package [Stuart et al., 2019] was used to visualize nuclei using tSNE (Fig. 2B,E), to perform nuclei clustering (Fig. 2F; Fig. S16), to infer and plot marker genes of nuclei clusters (Fig. S15). For nuclei clustering, resolution parameters 0.095 (CN), 0.04 (CB), and 0.15 (AC) were used. To assign cell type identity to clusters, we chose cell type marker genes based on literature: *GAD1, GAD2* (inhibitory neurons [Lake et al., 2016]); *SLC17A7, SATB2* (excitatory neurons [Lake et al., 2016]); *TAC1, PCDH8, DRD2, ADORA2A, PENK* (spindle neurons [Gokce et al., 2016; McCullough et al., 2018]); *PCP4, NECAB2, LMO7, CALB1* (Purkinje cells [Uhlen et al., 2015]); *PDGFRA, CSPG4* (oligodendrocyte precursor cells [Zeisel et al., 2018; McKenzie et al., 2018]); *GJA1* (astrocytes [McKenzie et al., 2018; Zeng et al., 2012]); *MBP, MOBP, MOG* (oligodendrocytes [Zeng et al., 2012]); *RELN* (cajal retzius [D’Arcangelo et al., 1997]); *AIF1, CX3CR1, PTPRC, HLA-DRA* (microglia [McKenzie et al., 2018]); *A2M* (endothelial vascular cells [Zeng et al., 2012]); *TIAM1* (granular cells [Uhlen et al., 2015]). We further plotted average expression levels of selected marker genes in each nucleus (Fig. 2F) and in each tSNE cluster (Fig. 2G).

### Human-specific expression differences in snRNA-seq dataset

Similar to RNA-seq data analysis, we classified human-specific expression differences in each cell type as the ones showing 2-fold greater human/macaque difference relative to chimpanzee/macaque or bonobo/macaque differences. Chimpanzee-specific and bonobo-specific expression differences were defined as the ones showing 2-fold greater chimpanzee/macaque or bonobo/macaque difference relative to human/macaque difference. To balance the number of nuclei between cell types, we bootstrapped the nuclei 1,000 times to 50 nuclei per cell type per region for this analysis. To assess the evolutionary rate of particular cell type, we first calculated the average number of human-specific, chimpanzee-specific, and bonobo-specific genes among 1,000 bootstraps for each cell type in three brain regions. Then, to facilitate comparison between regions, we divided the evolutionary rates by their mean in each region (Fig. 3A,B). Further, the human-specificity ratio of each cell type was estimated as the number of human-specific genes divided by the number of chimpanzee-specific or bonobo-specific genes (Fig. 3C,D). To assess heterogeneity of nuclei within clusters, we re-classified human-specific genes while bootstrapping the nuclei 1,000 times to one nucleus per cell type per region. Then, we calculated the pairwise overlap/union of human-specific genes among 1,000 bootstraps within each cell type and each region (Fig. S17).

We further averaged gene expression values per cell type per region in snRNA-seq data. Based on these values, we calculated the average human-specificity per cell type per region as the mean of log-transformed ratios of human/macaque difference to chimpanzee/macaque or bonobo/macaque difference in snRNA-seq data. The number of nuclei per cell type per region was balanced to 50 nuclei per cell type per region for this analysis. Pearson correlation coefficients were calculated between average gene expression values in human (Fig. 3E), and between average human-specificity per cell type per region (Fig. 3F).

To deconvolute bulk human-specific expression differences using a neuronal evolutionary signature, we first compared human-specificity ratios of genes preferentially expressed in neuronal subtypes (Table S5) between single nuclei and bulk RNA-seq datasets (Fig. 3G). For each of 33 brain regions, we further calculated the number of genes showing human-specific expression in bulk RNA-seq dataset, which overlapped with genes showing human-specific expression in at least one of the neuronal subtypes in snRNA-seq data (Fig. 3H,I).

### Gene expression differences detected by snRNA-seq and bulk RNA-seq

Expression differences separating humans from chimpanzees and bonobos in bulk RNA-seq dataset were defined as > 2-fold difference in human samples compared to a pool of chimpanzee and bonobo samples, BH-adjusted p < 0.05, two-sided t-test. For this analysis, we normalized log-transformed read counts per gene by the sample median but did not normalize by the median expression level among regions in each individual brain, for consistency with snRNA-seq dataset.

In snRNA-seq dataset, expression differences separating humans from chimpanzees and bonobos were defined in each cell type as > 2-fold difference in human nuclei compared to a pool of chimpanzee and bonobo nuclei, BH-adjusted p < 0.05, Wilcoxon test implemented in *Seurat 3*.*0* function FindMarkers [Stuart et al., 2019]. Only genes that were detected in a minimum fraction of 0.1 nuclei in humans or in a pool of chimpanzees and bonobos were tested.

To find an overlap between gene expression differences detected by snRNA-seq and bulk RNA-seq, we calculated the percentage of differences detected by both snRNA-seq and bulk RNA-seq relative to the total number of differences detected by bulk RNA-seq (Fig. 4A,B). Cell type-specific differences were solely detected in one cell type, while shared differences were detected in > 1 cell type by snRNA-seq (Fig. 4B). We further calculated a log10-transformed amplitude of these cell type-specific and shared expression differences measured using bulk RNA-seq between humans and the average of two ape species (Fig. 4C).

Next, we focused on gene expression differences detected by snRNA-seq but not by RNA-seq (Fig. 4D,E). Functional enrichment analysis was performed for each group of cell type-specific and shared differences using *clusterProfiler* (enrichGO function, BP ontology, genes expressed in AC snRNA-seq dataset as a background, BH corrected p < 0.01; Fig. S18) [Yu et al., 2012]. We calculated the percentage of cell type-specific and shared differences relative to the total number of differences solely detected by snRNA-seq (Fig. 4F). As there was an unequal number of nuclei per cell type, we observed more significant expression differences separating humans from chimpanzees and bonobos for cell types containing more nuclei because of the higher statistical power due to the larger sample sizes. Thus, for the unbiased percentage calculation, we balanced the number of nuclei per cell type to 110 human nuclei and a pool of 110 chimpanzee and bonobo nuclei by subsampling the nuclei 20 times and counting expression differences separating humans from chimpanzees and bonobos in ≥ 0.9 subsampling iterations.

### Immunohistochemistry image processing

AC sections immunostained with antibodies against NFAT5 and MSI2 proteins were subjected to quantitative analysis. Astrocytic processes density was calculated in three sections per sample in three humans, three chimpanzees, and three macaques (Fig. 4I; Fig. S19). To track expression inhomogeneity among cortical layers, tiles consisting of 2×6 fields of view (Fig. 4L) were stitched to cover upper part (∼1 mm) of the cortex using ZEN (Zeiss). Intellesis ZEN Module was used to segment NFAT5- or MSI2-positive astrocytic processes from neuronal nuclei and background. Mean density of segmented objects was measured for each image (Fig. 4J; Fig. S19). To test the significance of differences between species and between cortical layers in each of the species, two-sided t-test with Holm-Sidak correction was performed.

## DATA ACEESS

All raw and processed sequencing data generated in this study have been submitted to the NCBI Gene Expression Omnibus (GEO; http://www.ncbi.nlm.nih.gov/geo/) under accession numbers GSE127898 and GSE127774. The analysis software developed for this paper is available at https://cb.skoltech.ru/~khrameeva/brainmap/code/. We provide an interactive website at https://nucseq.cobrain.io/, reproducing key analyses of RNA-seq and snRNA-seq data with variable parameters. The website can be browsed by gene, with information on its expression level in cell types of four species.

## ACKNOWLEDGMENTS

The authors thank the Chinese Brain Bank Center for providing the human samples, and the Southwest National Primate Research Center, Lola Ya Bonobo Sanctuary, and the Suzhou Experimental Animal Center for providing the chimpanzee, bonobo, and macaque samples. We are grateful to CoBrain IT team and personally to Dmitry Vinogradov for the technical support. We thank Grace Cuddihy for her comments on the manuscript. This work was supported by the Strategic Priority Research Program of the Chinese Academy of Sciences (grant XDB13010200); the National Natural Science Foundation of China (grant 31420103920); the National One Thousand Foreign Experts Plan (grant WQ20123100078); the National Key R&D Program of China (grant 2017YFA0505700); and the Russian Science Foundation (grant 16-14-00220).

## AUTHOR CONTRIBUTIONS

B.T., J.G.C., S.P., C.S., and P.K. designed the research. D.H., P.G.L., S.K., M.S., and Z.Q. performed research. E.K., I.K., S.R., P.M., and A.T. analyzed the data. All authors contributed to writing the paper.

